# RNAvigate: Efficient exploration of RNA chemical probing datasets

**DOI:** 10.1101/2023.04.25.538311

**Authors:** Patrick S. Irving, Kevin M. Weeks

## Abstract

Chemical probing technologies enable high-throughput examination of diverse structural features of RNA including local nucleotide flexibility, RNA secondary structure, protein- and ligand-binding, through-space interaction networks, and multi-state structural ensembles. Performing these experiments, by themselves, does not directly lead to biological insight. Instead, deep understanding of RNA structure-function relationships typically requires evaluating a system under structure- and function-altering conditions, linking these data with additional information, and visualizing multi-layered relationships. Current platforms lack the broad accessibility, flexibility, and efficiency needed to iterate on integrative analyses of these diverse, complex data. Here, we share the RNA visualization and graphical analysis toolset RNAvigate, a straightforward and flexible Python library. RNAvigate currently automatically parses twenty-one standard file formats (primary sequence annotations, per- and inter-nucleotide data, and secondary and tertiary structures) and outputs eighteen plot types. These features enable efficient exploration of nuanced relationships between chemical probing data, RNA structure, and motif annotations across multiple experimental samples. Compatibility with Jupyter Notebooks enables non-burdensome, reproducible, transparent and organized sharing of multi-step analyses and data visualization strategies. RNAvigate simplifies examination of multi-layered RNA structure information and accelerates discovery and characterization of RNA-centric functions in biology.

## Introduction

Primary, secondary, tertiary, and quaternary structural features all affect RNA function. Primary structure, the nucleotide sequence of an RNA transcript, is determined by transcription start and termination sites, splice sites, and polyadenylation sites. RNAs can contain diverse post-transcriptional modifications, which expand the primary sequence alphabet. Secondary structure is the pattern of both canonical and non-canonical base pairing and can involve stable helical regions, pseudoknots, and conformationally flexible regions. Tertiary structure, the three-dimensional arrangement of atoms in an RNA transcript involves through-space interactions between nucleotide groups and can be stabilized by binding of ions, small molecules, or macromolecules. Quaternary structure, the multi-molecular structure of an RNA, includes its protein and RNA partner interactions.

These diverse levels of RNA structure can be interrogated with a large and expanding high-throughput toolbox (Table S1). Primary sequence features can be determined by sequencing-based methods that measure transcript complexity (1) and numerous post-transcriptional modifications (2). Computational methods that model RNA secondary structure can (and generally should) be augmented by incorporating data from structure probing experiments.

Current, high-information structure probing methods employ structure-specific chemical probes that form covalent adducts with RNA and are widely used to characterize and model RNA secondary structure. Chemical adducts can be encoded into complementary DNA (cDNA) via reverse transcription (RT) as either mutations (mutational profiling or MaP) or RT stops and quantified, per-nucleotide, using high-throughput sequencing (3, 4). Single-molecule correlated chemical probing (smCCP) strategies take advantage of the ability of MaP technology to detect multiple (potentially correlated) chemical events per molecule and provide deep insights into RNA secondary and tertiary structure, coordinated networks of protein-binding, and conformational ensembles (5). Proximity crosslinking methods use photo or chemical crosslinking to characterize RNA inter- and intramolecular interactions and RNA-protein binding (6–8). Both experimental and computational methods are becoming powerful tools for modeling higher-order RNA structure and macromolecular complexes (9–11).

These strategies produce rich and complex starting data that report on interdependent structural elements in RNA. These datasets also pose two broad challenges for data exploration and hypothesis generation. First, it is typically important to explore data filtering and preprocessing steps, such as background-correction and normalization, to make decisions on how best to process and interpret raw data. Second, many studies require that multiple layers of data be visualized, in combination and as a function of experimental conditions. Many solutions exist for examining individual aspects of RNA structure. Missing has been a flexible and easy-to-use data analysis toolset that enables integration of multiple complex datasets to produce accessible visualizations, in the multiple formats commonly used by the RNA community, and that support efficient biological hypothesis generation.

Jupyter notebooks are open-source interactive computing environments that work in modern web browsers and integrated development environments (12). These notebooks enable melding of analyses, visualizations, and observational notes. A single document thus creates an ideal environment for data exploration and hypothesis generation, a documented report of analysis findings, and a shareable and reproducible analysis pipeline. The widespread use and popularity of Jupyter notebooks has resulted in a well-developed ecosystem of open-source tools supporting collaboration on and publication of data analyses. However, until now, creation of the types of plots used by the RNA community required fluency in one of the programming languages compatible with Jupyter Notebooks.

Here we share a Jupyter-compatible Python module, RNAvigate, which broadly simplifies creation of RNA community-standard visualizations and analyses derived from diverse data sources. RNAvigate rapidly aligns sequences then compares, filters, analyzes, and displays RNA structural information. These features significantly streamline the analysis of complex datasets and the generation of impactful hypotheses regarding RNA structure-function interrelationships.

## Results and Discussion

### RNAvigate scope

RNAvigate is a highly flexible data visualization and communication tool. Initial processing of structure probing data is performed by more specialized software, for example, ShapeMapper (13, 14) to analyze per-nucleotide reactivities; or DanceMapper, DREEM or DRACO (15–17) to analyze single molecule correlated chemical probing (smCCP) experiments. The raw information provided by these programs, by itself, is typically insufficient to understand a particular RNA system deeply. Indeed, the work of and thinking about large-scale chemical probing data has just begun.

Individual chemical probing experiments are most advantageously interpreted in context with additional information, including measures of global structure, models of secondary and tertiary structure for individual RNA motifs, estimates of model confidence, and relationships to other RNA-centric features, including regulatory elements and RNA-binding proteins. RNAvigate accepts initial information from a wide variety of state-of-the-art technologies, is extensible to many more, and then facilitates integrative analysis of multiple classes of RNA-centric information and modeling.

We find that students who have never programmed or are new to the command line quickly learn to use RNAvigate to understand their data deeply and to produce complex visualizations that would otherwise be inaccessible to them. RNAvigate efficiently generates the complex illustrations that have proven especially impactful within the RNA structure community in generating mechanistic models and biological hypotheses.

### RNAvigate workflow

The RNAvigate workflow is organized into three steps (Figure 1). First, Inputs are curated from the raw output of experiments, computations, and database searches. Second, Data classes are created by providing RNAvigate with the input file names. Finally, visualizations are created using plotting functions built into RNAvigate.

**Figure 1.**
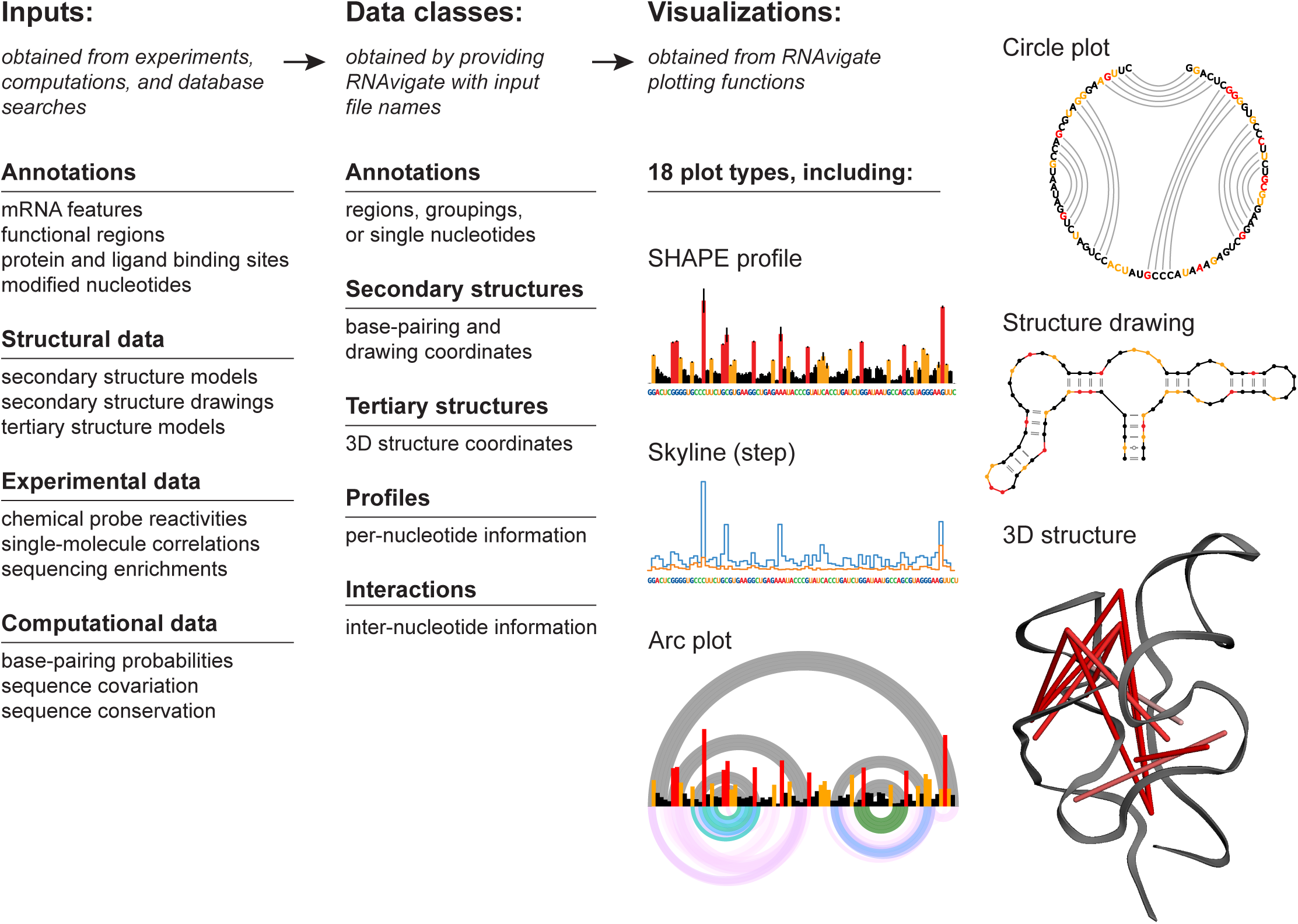
Workflow for data exploration using RNAvigate.

RNAvigate includes a high-level interface, presented here, in which the second and third steps in this workflow are each executed by a single command. These commands are designed to be simple to use, to automate many otherwise tedious tasks, and to remain highly flexible. This design choice allows users to access RNAvigate features without learning object-oriented programming. For users interested in the low-level, object-oriented underpinnings of RNAvigate, the full API is documented online (see Code Availability statement). This two-tiered design makes RNAvigate a powerful tool for both RNA bench scientists (including those with limited programming experience) and software and data specialists.

### Inputs and file formats

RNAvigate automatically parses RNA structural information in many standard file and data formats. The simplest inputs are annotations of the primary sequence of RNA. These annotations describe transcript features (translation start sites, exon junctions, UTRs), functional regions, modified nucleotides, or binding sites for proteins, ligands, and other nucleic acids.

RNAvigate also accepts file formats output by structure modeling software (including RNAStructure and Vienna RNA (18, 19)) and secondary structure drawing software (XRNA, VARNA, StructureEditor, FORNA, and R2DT (18, 20–22)). Supported tertiary structure formats include CIF and PDB. Inputted experimental data can include chemical probe reactivities, such as those from MaP experiments, single-molecule events detected by smCCP, and sequencing enrichments such as from CLIP experiments. Inputted computational data can include pairing probabilities, covariation, and sequence conservation. Most widely used file formats are natively supported by RNAvigate, and additional formats can be supported with simple code additions (Table S1).

RNAvigate is primarily designed for analyzing single transcripts. However, RNAvigate can also read genome sequences (fasta), transcript annotations (GFF, GTF), and genomic or transcriptomic per-nucleotide or region data (BED/NarrowPeak or WIG). If provided with transcript identifiers, such as ENSEMBL or RefSeq IDs, RNAvigate extracts data in transcript coordinates. This information can include sequence, exon junctions, coding and untranslated regions, per-nucleotide measurements, protein-binding sites (eCLIP), and other annotations.

### Data classes

The diverse sources of input data described above are stored and represented in RNAvigate as one of five data classes: annotations, secondary structures, tertiary structures, profiles, or interactions (Figure 1) These classes create functionality for the data to be analyzed, filtered, and visualized. Input data files are converted to one of these data classes when assigned to an RNAvigate data keyword. This assignment also organizes data in a standardized way across experimental samples and simplifies downstream commands.

Annotations contain regions of interest, single and grouped nucleotides of interest, and primer-binding sites. Secondary structures contain base-pairing information and optional diagram-drawing coordinates for secondary structure models. Tertiary structures contain atomic coordinates for tertiary structure models. Profiles contain any type of experimental or computational measurement made on a *per-nucleotide* basis. Examples of per-nucleotide data include chemical probing reactivities, structure-based entropies, and sequence conservation. Interactions contain experimental or computational measurements made *between* two nucleotides. These measurements include sequence covariation, base-pairing probabilities, single-molecule correlations, and crosslinking information. These five classes of transcript-centric information are flexible and extensible and allow RNAvigate to represent most types of RNA structural data (Table S1).

### Visualization options

RNAvigate automates many community-standard analyses and visualizations with easy-to-use, flexible functions. Two standard visualizations commonly used by the RNA community are per-nucleotide graphs and connectivity diagrams. Per-nucleotide graphs display nucleotide position along the x-axis and a measured value on the y-axis (Figures 1 and 2B). RNAvigate can output two types of per-nucleotide graphs: colored bar graphs for displaying a single dataset and stepped-line graphs (skyline plots) for comparing multiple datasets (Figure 1).

**Figure 2.**
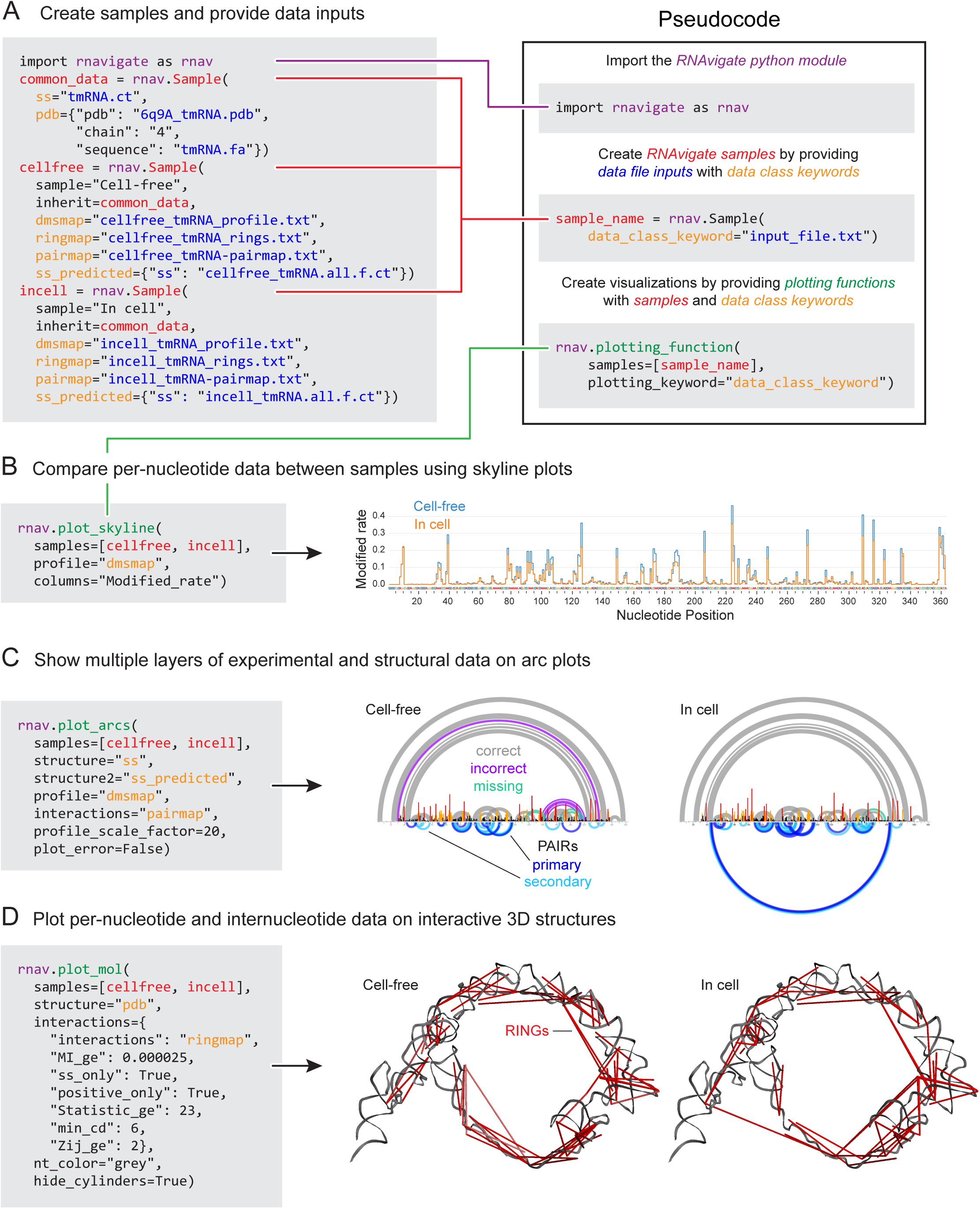
A complete RNAvigate workflow: single-molecule correlated chemical probing (smCCP) of tmRNA. In this example, full RNAvigate code is shown for each figure. (*box*) Essential coding steps in an RNAvigate analysis explained with code-like examples. (A) Importing RNAvigate module and loading in-cell and cell-free DMS probing data. These steps create a Python object that contains all data in a structured format. (B) Skyline plot comparing DMS-induced in-cell and cell-free mutation rates per nucleotide. (C) Arc plots illustrating multiple layers of experimental and structural data including literature-accepted structure, data-informed minimum free energy structure predictions, and PAIRs. Base-pairs are labelled “correct” (grey) if predicted and in the accepted structure, “incorrect” (purple) if predicted and not in the accepted structure, and “missing” (green) if not predicted and in the accepted structure. PAIRs of the highest confidence are labelled “primary” (dark blue), otherwise are “secondary” (light blue). (D) Visualization of three-dimensional tmRNA backbone (gray), overlayed with cell-free and in-cell RINGs (red). Data are from ref. (27).

RNAvigate can create four types of connectivity diagrams: arc plots, circle plots, secondary structure diagrams, and interactive three-dimensional molecular renderings (Figures 1, 2C-D, 3, 4A-C, 5A-C, and 6C). Connectivity diagrams display through-space intra-nucleotide interactions such as base-pairs, crosslinks, or correlated single-molecule events. Arc plots arrange nucleotides along the x-axis and show interactions as semi-circles. Circle plots arrange nucleotides around a circle with interactions drawn as parabolas. Secondary structure diagrams have custom arrangements of nucleotides and display interactions as lines. Finally, interactive three-dimensional molecule renderings arrange nucleotides according to atomic coordinates and display interactions as thin cylinders (Figure 1).

Other visualizations automated by RNAvigate include 3D-distance histograms, heatmaps (Figure 4B), and contour plots (8, 23). Automated analyses include identification of regions with low SHAPE reactivity and low Shannon entropy (lowSS regions) (24) (Figure 2), linear regressions (Figure 6A), deltaSHAPE (25), and windowed area under receiver-operator characteristic curves (26).

### Example workflows

The following sections provide examples for using RNAvigate to analyze RNA structure and to generate hypotheses regarding underlying RNA-mediated function. Figures 3-5 replicate published figures and illustrate the ability of RNAvigate to create high-content illustrations efficiently. Figure 6 compares data from four independent studies and illustrates the flexibility of RNAvigate in integrating analysis of data from multiple sources, in different formats. The raw data, full code, and Jupyter notebooks used to generate these analyses and figures are fully and freely available via GitHub and Binder, the latter allows interaction with these notebooks with no required software download or installation (see Methods and Data statement).

**Figure 3.**
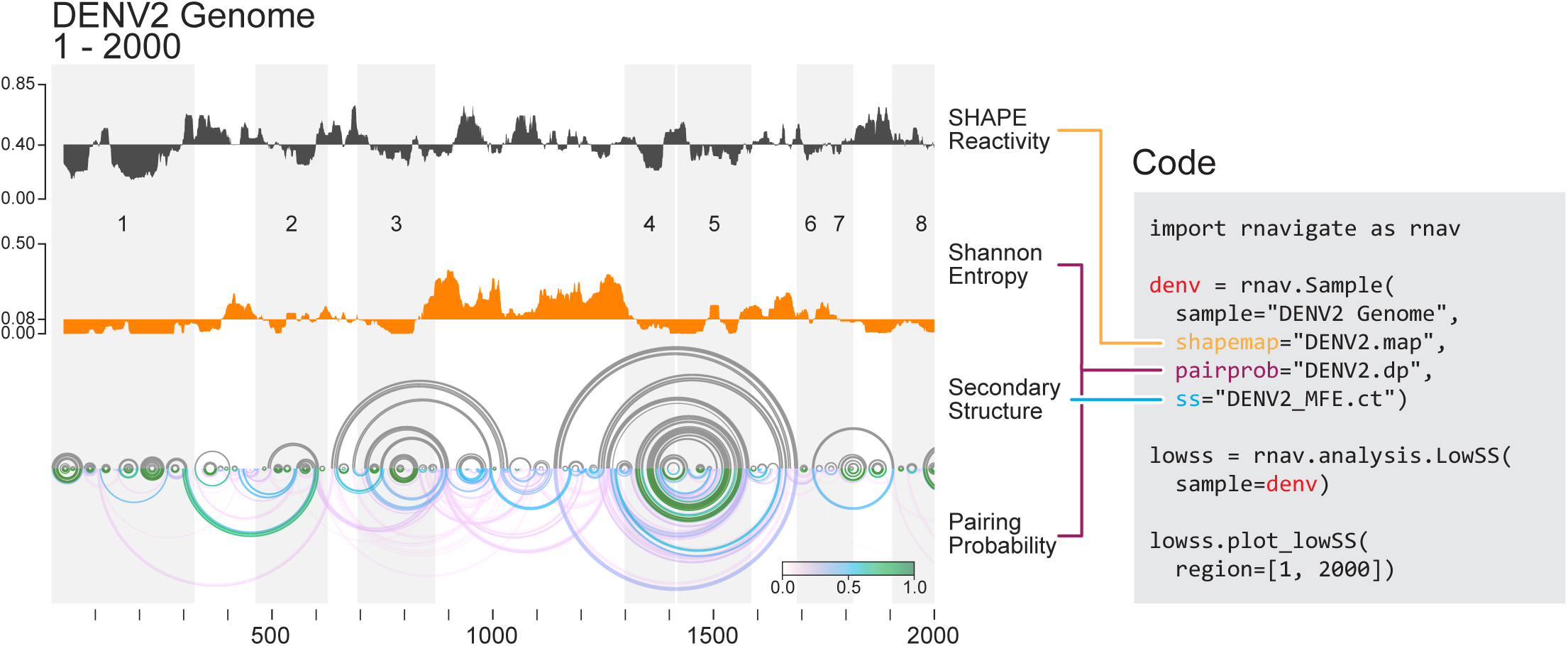
Identification and visualization of low-SHAPE–low-Shannon entropy (lowSS) regions in the DENV2 genome using RNAvigate. Median (*top*) SHAPE reactivity and (*middle*) Shannon entropies over 51-nucleotide windows across the DENV2 genome. (*bottom*) Modeled minimum free energy structure (gray) and pairing probabilities (see scale) illustrated as arc plots. Regions with median SHAPE reactivity below 0.4 and median Shannon entropy below 0.15 are defined as lowSS regions and are shaded in gray and numbered. Full RNAvigate code used to generate this analysis and visualization shown at right; data are from ref. (24).

### smCCP analysis

Single molecule correlated chemical probing (smCCP) strategies represent a revolution in RNA structure probing and allow multiple classes of through-space interactions to be measured in a simple chemical probing experiment (5). The MaP strategy allows multiple chemical adducts to be measured across the same RNA strand. When these chemical modification events occur at two nucleotides in *correlated* way, smCCP measures structural communication between these positions (5). smCCP experiments can be analyzed to directly observe, rather than merely infer, base pairing (pairing ascertained from interacting RNA strands or PAIRs), to measure through-space interactions reflective of tertiary structure (RNA interacting groups or RINGs), and to deconvolute RNA conformational ensembles. Whereas smCCP experiments are generally straightforward to perform, their analysis and interpretation can be complex.

RNAvigate dramatically streamlines the required analysis. We will illustrate one such analysis in a detailed way in this first example, showing all RNAvigate code explicitly. Similar use principles then apply to all examples shared here.

Bacterial transfer-messenger RNA (tmRNA) is responsible for rescuing stalled ribosomes. This RNA has a complex secondary structure, including four pseudoknots, and a large open ring tertiary structure. Secondary structures for tmRNAs are poorly predicted by computational methods alone (27). We used RNAvigate to facilitate analysis of publicly available structure probing data for in-cell and cell-free dimethyl sulfate (DMS) treated *E. coli* RNA (27). The original data set was used to assess the ability of PAIRs to improve modelling of RNA secondary structure in cells. We confirmed that result, and further explored through-space tertiary interactions measurable from these data, but not previously reported.

We collected raw sequencing reads from a sequence read archive, the literature-accepted secondary structure in the form of a .ct file, and the cryo-EM structure (28) from the RCSB. Raw sequencing reads were then analyzed with ShapeMapper2, PairMapper and RingMapper (13, 27). ShapeMapper2 calculates per-nucleotide mutation rates reflective of DMS reactivity with RNA. PairMapper detects smCCP events indicative of base-pairing (PAIRs). RingMapper detects smCCP events arising from through-space interactions (RINGs). DMS reactivities and PAIRs were then used to direct minimum free energy-based secondary structure modeling using the RNAstructure fold program (18) (using -dmsnt and -x parameters, respectively).

Within a Jupyter notebook, the RNAvigate Python module was imported and given the alias **rnav** (Figure 2A). Input file names were provided for each sample using **rnav.Sample()**. This important step accomplishes three things: the data contained in each file are associated with an experimental sample, converted into a data class, and assigned to a short keyword for easy access. Data for cell-free and in-cell experimental samples were assigned to variable names **cellfree** and **incell**, respectively (Figure 2A). We illustrate here three classes of smCCP data analysis for the tmRNA; each plot is created with a single command, optionally with modifying arguments (Figure 2B-D).

We first visualized overall reactivity profiles under in-cell and cell-free conditions, using a skyline plot (Figure 2B). Per-nucleotide reactivity rates were highly similar overall, and slightly higher under cell-free conditions, reflective of higher structure for this non-coding RNA in cells, a common observation. Next, an arc plot was used to compare modeled and accepted structures and to visualize the DMS reactivities and PAIRs that informed the structure model. This visualization revealed that in-cell data produced a more accurate model of the secondary structure than did data obtained under cell-free conditions. The RNAvigate analysis showed that increased accuracy (relative to the functional structure) is driven by more abundant and accurate PAIRs for the in-cell data, as the per-nucleotide reactivity profiles are very similar for the two conditions (Figure 2C). Finally, RINGs were visualized superimposed on the cryo-EM structure. These data reveal extensive structural communication between the adjacent pseudoknots along the outer ring of the structure and an absence of structural communication across the circle-like opening (Figure 2D).

### Analysis of regions of low SHAPE reactivity and low Shannon entropy

One especially informative analysis of long RNAs is to identify regions that have both low SHAPE reactivities, indicative of extensive base-pairing, and low Shannon entropy, which indicates a well-defined structure. These low SHAPE-low Shannon entropy (lowSS) regions are strongly associated with functional regions in long RNAs (24, 29, 30). RNAvigate makes identification and visualization of lowSS regions straightforward. Here, we use RNAvigate to reproduce a useful analysis based on a publicly available dataset obtained from SHAPE probing of Dengue virus serotype 2 (DENV2) genomic RNA (24).

Dengue virus has a positive-strand single-stranded RNA genome of about 11 kilobases. Conserved structures in the 5’ and 3’ untranslated regions regulate viral processes and are well characterized (31). In contrast, identification of functional structures in less conserved regions has presented challenges. Analysis of lowSS regions made it possible to define novel regions of functional importance and to identify regions likely to have complex higher-order (tertiary) structures (24, 32).

RNAstructure programs fold and partition were used to model a minimum free energy structure and determine base-pairing probabilities, respectively, using SHAPE reactivity data as pseudo-free-energy constraints (--shape parameter) (18). These results and RNAvigate were imported into a Jupyter Notebook. RNAvigate was used to calculate 51-nucleotide windowed median SHAPE reactivities and median Shannon entropies based on the provided pairing probabilities (Figure 3, *top* and *middle*). The program then identified lowSS regions where the windowed median SHAPE reactivity was below 0.4 and the median Shannon entropy was below 0.15.

RNAvigate enables visualization of this analysis as stacked profiles of SHAPE reactivity and Shannon entropy, and of predicted minimum free energy structures and pairing probabilities displayed as arc plots. lowSS regions are highlighted and numbered (Figure 3); plots share the same x-axis (nucleotide position) for easy comparison. Visualization of the first 2000 nucleotides of the DENV2 genome revealed eight lowSS regions.

This straightforward application of RNAvigate recapitulates a published analysis (24) that required significantly greater effort to create. The original illustration was created using command-line tools to generate a PDF image of the arc plot and a spreadsheet program to calculate and to visualize windowed SHAPE and Shannon entropy profiles. These plots were copied into a vector drawing program (Adobe Illustrator), where they were manually adjusted to align the x-axes and to highlight lowSS regions. Plot labels, legends, and axes were added manually. RNAvigate accomplishes a similar, if not improved, result in 3 lines of code; similar efficiencies are achieved with all of the following examples.

### Analysis of time-resolved probing

Many dynamic RNA processes are best analyzed in a time-resolved way, which creates complex datasets across multiple experimental samples. In one recent example, we developed a fast-reacting alkylating reagent, trimethyloxonium (TMO), and used it to follow the time-resolved folding of the catalytic core of a ribonuclease P (RNase P) enzyme (33). The folding pathway was examined in smCCP experiments and involved creating multiple secondary and tertiary structure diagrams and heatmaps of through-space RINGs. This complex analysis is now rendered straightforward with RNAvigate.

*B. stearothermophilus* RNase P is a long non-coding RNA that catalyzes cleavage of 5’ leader sequences of precursor tRNAs, an essential step for tRNA maturation. TMO reacts with RNA via a chemical mechanism similar to that of DMS but reacts 90 times faster. TMO was used to perform time-resolved probing of RNase P, and smCCP analysis revealed the through-space interaction networks that form and change during the folding of this large RNA (33). The in vitro-transcribed RNase P RNA was treated with TMO, initially in the absence of Mg^2+^, and then at multiple time points after adding Mg^2+^. Raw sequencing reads corresponding to MaP analyses at each time point and a custom-drawn secondary structure were obtained from the original publication. The raw data were analyzed using ShapeMapper2 and RingMapper. A three-dimensional model of the RNA (based on PDB 3DHS (34)) was obtained from the authors. RNAvigate was used to import the resulting files into a Jupyter notebook.

For the RNAvigate analysis, a detailed secondary structure was generated to orient the reader to the functional domains, important interactions, and helices present in RNase P, particularly the catalytic core, the P2-P5 pseudoknot, and the L5.1-L15.1 loop-loop interaction (Figure 4A). Next, the strongest RINGs with positive correlations were displayed on the secondary structure drawing. This superposition revealed large changes in the structural communication network as folding progresses. The L5.1-L15.1 loop-loop interaction forms very quickly and is present in an early partially folded state. In the fully folded state, interactions within the catalytic core are more pronounced, revealing this structure forms more slowly (Figure 4B, *top*).

**Figure 4.**
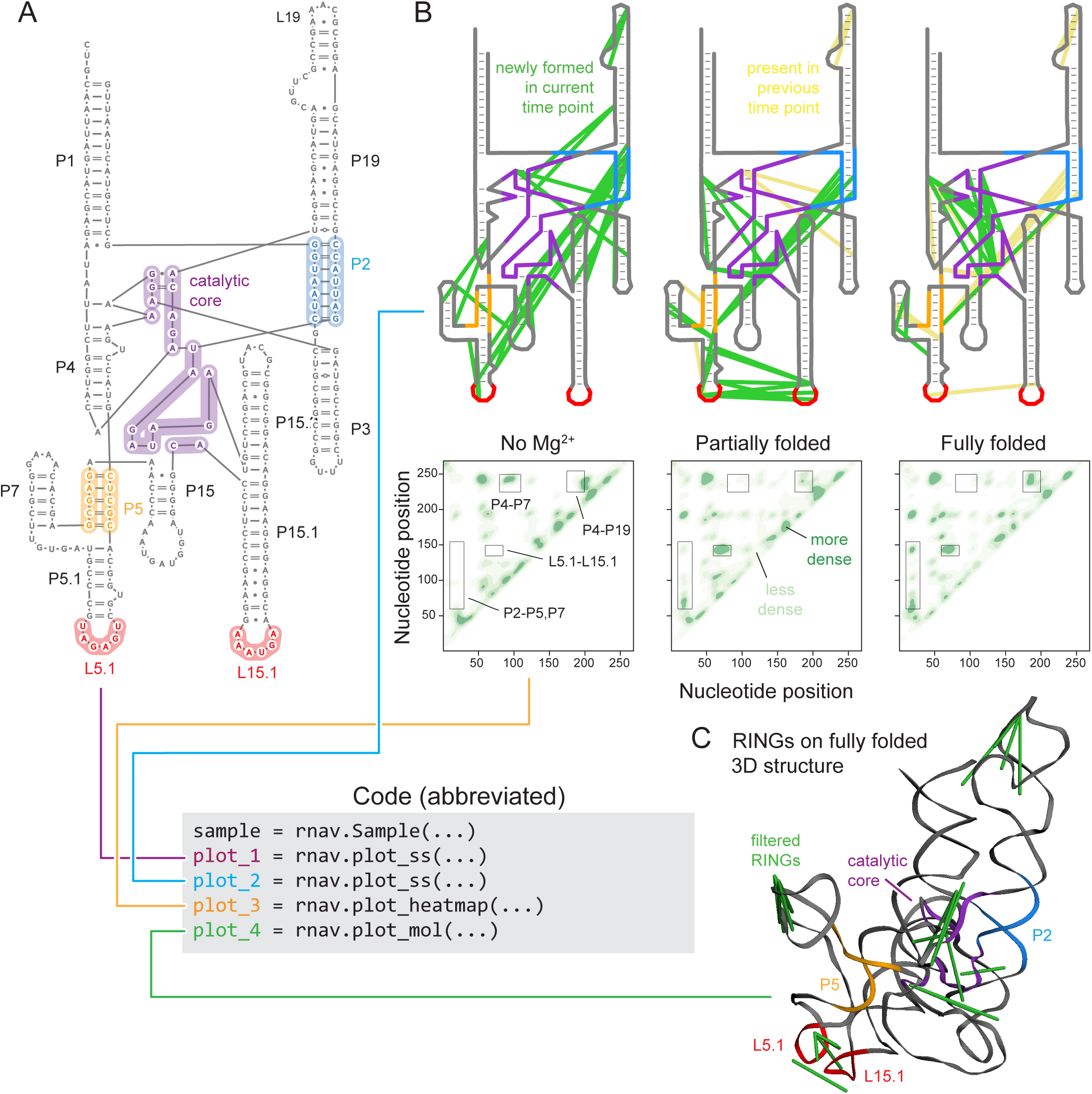
Time-resolved probing and folding of the RNase P RNA, visualized using RNAvigate. (A) Secondary structure of the catalytic core of RNase P RNA. Regions involved in major time-dependent changes are highlighted. (B) (*top*) Through-space RINGs as a function of folding time superimposed on the secondary structure. Newly formed RINGs at each timepoint are shown in green; RINGs retained from the prior timepoint are yellow. (*bottom*) Heatmaps of RING density at each time point. (C) Three-dimensional structure of the RNase P RNA with superimposed RINGs (in green). Data are from ref. (33).

The spatial density of positively correlated RINGs, plotted as heatmaps, across the three states revealed additional interactions where through-space RING correlations change during the folding process (Figure 4B, *bottom*). This visualization, like the RINGs displayed on the secondary structure above, identified rapid formation of the L5.1-L15.1 loop-loop interaction upon addition of Mg^2+^. These heatmaps also reveal that interactions in the P4-P7 region disappear and interactions in P2-P5,P7 region form quickly after addition of Mg^2+^, indicating interdependency between the pseudoknot helices. RING densities also reveal a slower-forming interaction between P4 and P19. Finally, RINGs from the fully folded state were filtered to isolate likely tertiary interactions and plotted on the three-dimensional structure. These RINGs illustrate the density of interactions in the catalytic core and suggest this fully folded state closely matches the known structure (Figure 4C). Complex, time-resolved data are now analyzed efficiently using RNAvigate.

### Deconvolution of a conformational ensemble

smCCP strategies can now resolve complex conformational ensembles of RNA, including in the cellular environment (5). The DANCE (deconvolution and annotation of ribonucleic conformational ensembles) framework uses machine learning to deconvolute smCCP data from a DMS-MaP experiment to define the dominant states in an RNA conformational ensemble and can then assign through-space PAIRs and RINGs to each state. This single experiment and subsequent analyses produce multiple layers of data for each state including population percentages, reactivity profiles, PAIRs, RINGs, predicted secondary structures, and pairing probabilities (15). RNAvigate makes analysis of this layered information, and closely related smCCP experiments (7, 16, 17, 35), broadly accessible.

The adenine riboswitch region of the *V. vulnificus add* mRNA populates two dominant conformations. Adenine binds to the ON state and shifts the equilibrium toward this state. Formation of the ON state results in production of adenine deaminase because, in this conformation, the Shine-Dalgarno (SD) sequence of the mRNA is single-stranded and accessible to the ribosome. In the OFF state, the Shine-Dalgarno sequence is sequestered in a duplex. Conventional per-nucleotide structure probing methods measure population averages and are misleading for this riboswitch, because the RNA samples multiple well-populated conformations (15).

For RNAvigate analysis, data files from the DANCE-MaP analysis of the adenine riboswitch in the absence of adenine (15) were obtained from the GEO (GSE182552). These single-experiment data files contain reactivity profiles, PAIRs and RINGs for each of the dominant conformations. The foldClusters module of DanceMapper was used to predict minimum free energy structure models for each conformation, based on providing the reactivities (required input) and PAIR constraints (--bp parameter) for each conformation. These structure models were loaded into StructureEditor (18) and manually arranged to match the original publication. A three-dimensional structure for the translation ON state was obtained from the PDB (4TZX) (36). File names were then provided to RNAvigate within a Jupyter notebook and used to create the visualizations in this example.

As visualized using arc plots and consistent with the original publication (15), PAIRs accurately and sensitively detected most base-paired helices, recapitulating the accepted secondary structures of, and the differences between, these two states (Figure 5A, in *blue*). RINGs captured the loop-loop tertiary interaction stabilizing the three-dimensional structure of the riboswitch ON state (Figure 5B-C, in *red*). In the context of the crystal structure, through-space RING interactions link two close-in-space loops (Figure 5C). RNAvigate efficiently analyzes information-rich DANCE-MaP experiments and straightforwardly visualizes the multiple dominant states present in an RNA structural ensemble.

**Figure 5.**
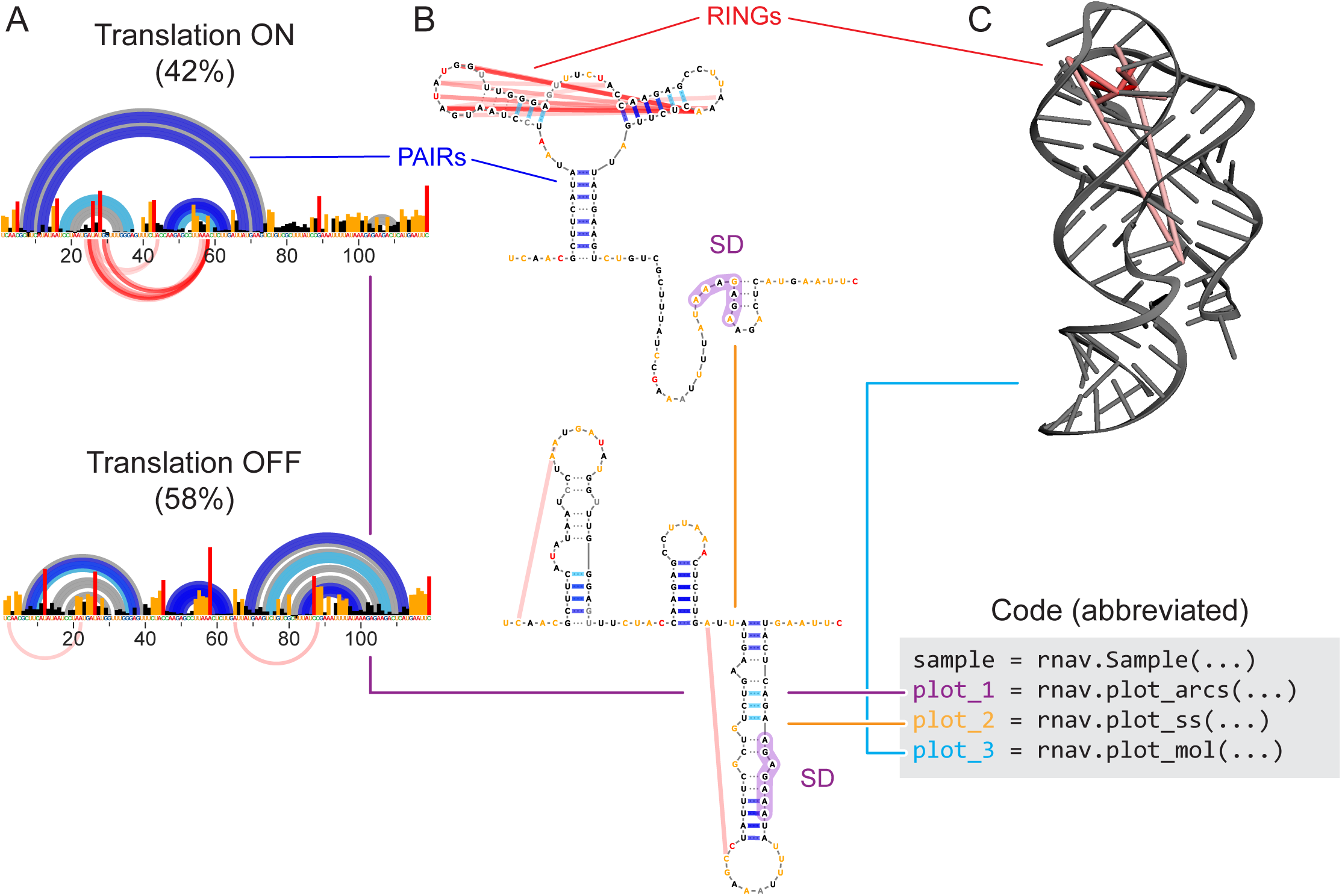
DANCE-MaP analysis of the *add* riboswitch ensemble, visualized using RNAvigate. (A) PAIRs, RINGs (each shown as arc plots), and reactivity profiles for DANCE-identified ON and OFF states. (B) Secondary structure drawings (conventional style) for each component, with PAIRs (blue) and RINGs (red) superimposed. Shine-Dalgarno sequence (SD) is highlighted. (C) Structure of *add* riboswitch in the ligand-bound state, overlayed with RINGs uniquely identified in this state. Note that these data were all obtained in the absence of adenine ligand but, nonetheless, clearly show that the ade riboswitch populates the ON state. Data are from ref. (15).

### Comparison of chemical probing methods

Comparing chemical probing datasets from wide-ranging experiments can be incisive for understanding differences in experimental choices or between functional and cellular states. Four groups recently reported secondary structure models of the SARS-CoV-2 RNA genome, based on in-cell structure probing (26, 37–39). Here we use RNAvigate to compare these per-nucleotide reactivities and secondary structure models, and to evaluate the overall robustness of chemical probing to produce consensus secondary structure models.

The SARS-CoV-2 betacoronavirus is encoded by a single-stranded, positive-strand RNA genome of about 29 kb. The four studies examined here all chemically probed native viral RNA in cells, encoded chemical adducts into cDNA libraries, and computed per-nucleotide reactivities. These data were then used to guide thermodynamic modelling of viral RNA secondary structure. Each group executed these steps in distinct ways, performing either SHAPE or DMS probing, reading out sites of adducts using either the MaP or an RT-STOP strategy, using different RT enzymes, and differing in experimental and computational parameters (summarized in Table S2) (26, 37–39).

Per-nucleotide reactivities and secondary structure model data files were downloaded from the supporting data for each publication. Downloaded data files were either in standard formats (.ct, dot-bracket, ShapeMapper2 profile.txt, or RNAframework XML), all natively supported by RNAvigate, or in Excel spreadsheets (loaded via the Python command Pandas.read_excel). We focused on population average reactivities and structure models, across a representative region spanning the first 7000 kb of ORF1ab.

Reactivity data were unintuitively dissimilar, as indicated by Pearson correlation coefficients (0.19 < *r* < 0.49) and visualized using kernel density plots (Figure 6A). Nevertheless, 61% of modeled base pairs were shared across all studies (Figure 6B). All four studies based their structure modeling strategy on the ΔG_SHAPE_ (or ΔG_DMS_) framework (4). In this pseudo-free-energy change strategy, structure probing reactivities are converted into ΔG bonuses and penalties for base-pairing and used to modify and improve thermodynamics-based calculation of high-probability structures (4, 18). The similarity in structural models reinforces the foundational finding that this structure modelling strategy is robust for diverse underlying data. Modeling large structures typically requires imposing a maximum pairing distance constraint to prevent over-modeling of long-range base-pairs, the optimal value of which is generally not known. The four studies chose different constraints, ranging from 300 to 600 nucleotides, limiting similarity in structure models, especially for longer-range interactions (Figure 6C). These differences can have important functional consequences (4, 5), and can be resolved by the PAIR experiment (Figures 2 & 5).

**Figure 6.**
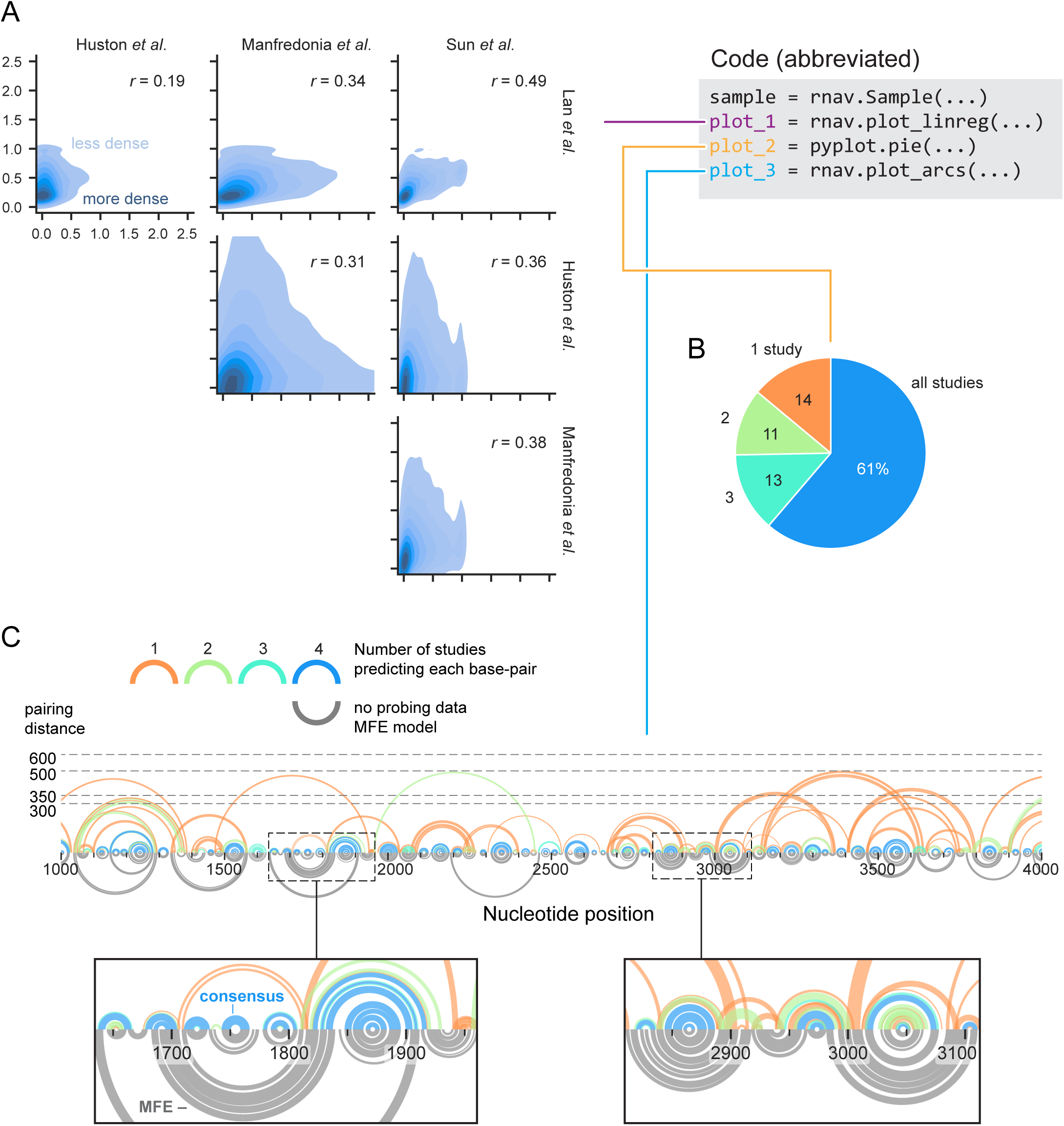
Comparative analysis of SARS-CoV-2 structure probing data and secondary structure models. (A) Study vs study per-nucleotide reactivities (shown as kernel density estimates) and Pearson correlation *r* from linear regression analysis. (B) Pie chart showing percentage of predicted base-pairs observed across studies. (C) Visualization of modeled base pairs as a function of study consensus (upper arcs), compared to a MFE structure (gray arcs). All predicted base pairs from all studies are shown (as arcs); colors indicate the number of studies predicting that base pair. Dotted lines indicate maximum pairing distance constraints, used by the different studies (modeled pairing arcs will not exceed this line). Analyses shown here correspond to 7000 kb of ORF1ab, where data were available for all studies (positions 266 through 7265). Data are from refs. (26, 37–39).

The data-driven models differed notably from MFE (minimum free energy, using no probing data) structure models. We found 24% of the 2,257 total base-pairs in a standard MFE model (calculated with max pairing distance = 300 (18)) do not appear in any probing-directed model, and 12% of the 1,234 consensus base-pairs (modelled in at least 3 studies) are not recovered. Differences between the data-driven and MFE models are not evenly distributed; some regions are modeled particularly poorly, and it is not possible to define *a priori* where a MFE model is most misleading (Figure 6C, *bottom*). In sum, RNAvigate efficiently compares reactivities and structure models from wide-ranging experimental strategies, identifies both consensus and divergent helices, and emphasizes there are many structural elements that are difficult to model without guiding per-nucleotide experimental information.

### Perspective

RNA structure-function interrelationships are complex. Powerful and orthogonal computational and experimental strategies have been developed to identify the hierarchical and interdependent elements of RNA structure. It is both important and challenging to understand, integrate, and interpret these rich layers of information. The visualization and analysis tools of RNAvigate facilitate analysis of diverse structural features of RNA. RNAvigate in conjunction with Jupyter Notebooks enable rapid analysis and thorough documentation of data quality control, data exploration, and hypothesis generation. RNAvigate then directly generates publication-quality figures.

RNAvigate, when used within a Jupyter notebook, provides a convenient, non-burdensome platform for documentation and for transparent and reproducible sharing, which enhances the longevity and impact of analyses of RNA experimental and computational data. Full pipelines for analysis and figure generation can be recreated, repurposed, or reconfigured using open-source web-based tools, without the requirement to download additional data or install specialized software, as demonstrated by the Jupyter notebooks accompanying this manuscript.

RNAvigate is modular and readily extensible, and thus provides a framework to support analysis and visualization of diverse experimental and computational data types. We anticipate that RNAvigate will enhance development, implementation, reproducibility, and sharing of wide-ranging RNA structure, interaction, and annotation strategies for discovery and characterization of diverse RNA-centric functions in biology.

## Methods and Data

Data files and Jupyter notebooks containing all data and methods described here are included in the supplemental data and are available on GitHub (https://github.com/Weeks-UNC/RNAvigate_figures). Notebooks on GitHub have been made interactive using the free Binder web service (part of the Jupyter Project). Users can reproduce these analyses and explore the data using RNAvigate in a web browser, with no required downloads or software installations.

## Code availability

The RNAvigate python module is available on Github (https://github.com/Weeks-UNC/RNAvigate) and is installable via Docker by following the instructions included in the documentation. RNAvigate documentation is available on Read the Docs (https://rnavigate.readthedocs.io).

## Acknowledgements

This work was supported by the NIH (R35 GM122532) and NSF (MCB-2027701) to K.M.W.

